# Reduced levels of membrane-bound alkaline phosphatase in Vip3Aa-resistant *Heliothis virescens*

**DOI:** 10.1101/2020.02.10.943167

**Authors:** Daniel Pinos, Maissa Chakroun, Anabel Millán-Leiva, Juan Luis Jurat-Fuentes, Denis J. Wright, Patricia Hernández-Martínez, Juan Ferré

## Abstract

The Vip3Aa insecticidal protein from *Bacillus thuringiensis* (Bt) is produced by specific transgenic corn and cotton varieties for efficient control of target lepidopteran pests. The main threat to this technology is the evolution of resistance in targeted insect pests, thus understanding the mechanistic basis of resistance is crucial to deploy the most appropriate strategies for resistance management. In this work, a laboratory-selected colony of *Heliothis virescens* (Vip-Sel) highly resistant to the Vip3Aa protein was used to test whether an alteration of membrane receptors in the insect midgut might explain the resistance phenotype. Binding of ^125^I-labeled Vip3Aa to brush border membrane vesicles (BBMV) from 3rd instar larvae from Vip-Sel was not significantly different from binding in the reference susceptible colony. Interestingly, BBMV from Vip-Sel larvae show dramatically reduced levels of alkaline phosphatase activity, which was further confirmed by a strong down-regulation of the membrane-bound alkaline phosphatase 1 (*HvmALP1*) gene. However, its involvement as a receptor for the Vip3Aa protein was not supported by ligand blotting and viability assays with insect cells expressing *HvmALP1*. These data support that reduced alkaline phosphatase, previously observed in insect colonies resistant to Cry proteins from Bt, may also serve as an indirect marker that is not mechanistically involved in resistance to Vip3Aa.

**IMPORTANCE:** The Vip3Aa insecticidal protein remains the only lepidopteran-specific trait in transgenic Bt crops with no cases of field-evolved resistance. While laboratory-selected resistance to Vip3A has been reported elsewhere, the mechanism for resistance is unknown. Results in this work show lack of significant Vip3Aa binding alterations in resistant and reference colonies of *H. virescens.* These observations are in contrast to most cases of high levels of resistance to insecticidal Bt proteins for which decreased binding is commonly detected. In addition, this study provides the first evidence of down-regulation of membrane bound alkaline phosphatase (mALP) associated with Vip3Aa resistance, a phenomenon commonly associated with resistance to Cry proteins from Bt. Results from this work suggest that mALP down-regulation may be a useful biomarker yet reject its direct participation in resistance to Vip3Aa.

## INTRODUCTION

The polyphagous pest *Heliothis virescens* (L.) (Lepidoptera: Noctuidae) is well known for producing substantial economic losses, particularly in cotton production, due to its ability to evolve resistance to different synthetic control products, such as methyl parathion or pyrethroids (1,2). As an alternative approach, genetically modified crops expressing insecticidal genes from *Bacillus thuringiensis* (Bt crops) were introduced in 1996 for the control of this and other pests. However, extensive use threatens their effectiveness and cases of field-evolved practical resistance have already been reported for some lepidopteran and coleopteran pests (3).

Gene pyramiding has been proposed as an effective strategy for insect resistance management in Bt crops (4). This approach consists of combined production of distinct insecticidal Bt proteins in the same plant, and its success heavily relies on the expressed proteins having distinct mode of action, commonly defined as not sharing binding sites in target tissues (5,6).

Although the mechanism of action and receptors for Cry proteins have been widely studied (7), little is known about the biochemical mechanisms that underlie the action of Vip3A proteins. Several studies have shown that Vip3A proteins do not share binding sites with Cry1 or Cry2 proteins, yet their damage to the midgut epithelium resembles Cry action (8-11). Supported by the lack of shared binding sites, transgenic corn and cotton varieties pyramided with Cry1, Cry2 and Vip3A genes are currently commercialized in several countries.

Knowledge of the biochemical and genetic level factors involved in resistance is crucial to design practices that delay the appearance of resistance or allow its rapid detection and ways to overcome it. The genetic potential to evolve resistance to Vip3A has already been shown in some laboratory-selected insect species such as *H. virescens* (12), *Spodoptera litura* (13), *Helicoverpa armigera* and *Helicoverpa punctigera* (14), and *Spodoptera frugiperda* (15,16). However, the biochemical basis of resistance to Vip3A has only been studied in a laboratory-selected colony of *H. armigera*, for which alteration of the binding sites was found not to be the cause of resistance (17).

In the present study, we aimed to determine the biochemical basis of resistance in a Vip3A-resistant *H. virescens* colony. In a previous study with this colony (Vip-Sel), resistance was shown to be polygenic (18), and a transcriptomic analysis showed significant differential gene expression with 420 over-expressed and 1,569 under-expressed genes (19). Results herein support that Vip3Aa binding is not altered significantly in resistant compared to susceptible *H. virescens*, and demonstrate an association between Vip3A resistance and down-regulation of a membrane bound alkaline phosphatase (mALP). Considering that orthologues of this mALP are commonly down-regulated in Cry1-resistant lepidopteran pests (20-23), our results suggest that the potential use of mALP down-regulation as resistance marker may extend to resistance against Vip3A proteins.

## RESULTS

### Vip3Aa binding to midgut brush border membrane vesicles (BBMV)

In testing whether binding of Vip3Aa was altered in the Vip3A-resistant (Vip-Sel) compared to the reference susceptible (Vip-Unsel) colony, we measured binding of Vip3Aa labeled with ^125^I to BBMV from the two colonies. Binding analyses showed specific Vip3Aa binding for BBMV from both colonies, with similar homologous competition curves (Figure 1, panel A). A high percentage of non-specific binding to the microtube walls was detected (35-40% of the input labeled toxin), which accounted for most of the non-specific binding (binding that is not blocked by high concentrations of unlabeled Vip3Aa competitor) observed in the competition curves. The *K*_d_ and *R*_t_ values estimated from the competition curves indicated that Vip3Aa binds with low affinity to a high number of binding sites in BBMV from *H. virescens* (Table 1). No significant differences (P > 0.05) were found in these equilibrium binding parameters between the two *H. virescens* colonies, suggesting that binding alteration is not mechanistically related to Vip3Aa resistance in Vip-Sel.

**Table 1.** Equilibrium *Kd* (dissociation constant) and *Rt* (concentration of binding sites) binding parameters estimated from Vip3Aa homologous competition assays with BBMV from resistant (Vip-Sel) and susceptible (Vip-Unsel) *H. virescens* insects.

| Strain | Mean ± SEM<sup>a</sup> | |
| --- | --- | --- |
|  | *Kd* (nM) | *Rt* (pmol/mg)<sup>b</sup> |
| Susceptible | 138 ± 18 | 443 ± 66 |
| Resistant | 161 ± 34 | 443 ± 109 |

<sup>a</sup> Values are the mean of two replicates.
<sup>b</sup> Values are expressed in picomoles per milligram of BBMV protein.

**Figure 1.**
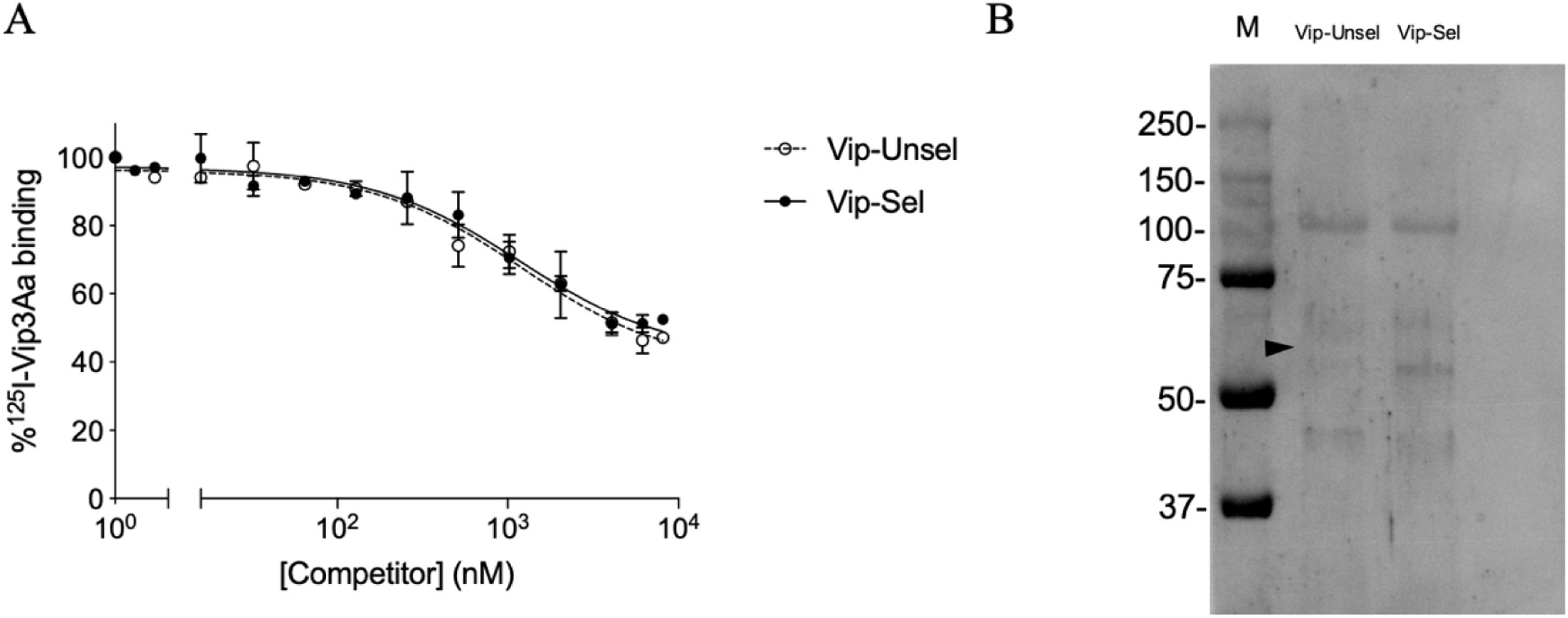
Analysis of the binding of BBMV from Vip-Unsel and Vip-Sel colonies of *H. virescens* to Vip3Aa. **A)** Competition binding assays of BBMV from the two colonies with ^125^I-Vip3Aa, using increasing concentrations of unlabeled Vip3Aa as competitor. Each data point represents the mean of two replicates (± SEM). **B)** Ligand blot performed with Vip3Aa against BBMV from Vip-Unsel and Vip-Sel colonies. Lane M: Protein marker (molecular weight in kDa). Black arrow indicates expected molecular weight of mALP (*ca.* 66 kDa).

### Reduced alkaline phosphatase (ALP) levels in the Vip-Sel colony

As a preliminary step to the use of brush border membrane vesicles (BBMV) in binding studies, we determined the specific activities of two brush border membrane markers, alkaline phosphatase (ALP) and aminopeptidase-N (APN), in larval midgut homogenates and BBMV preparations from susceptible (Vip-Unsel) and Vip3A-resistant (Vip-Sel) *H. virescens* colonies (Figure 2). The specific APN activity in midgut homogenates from both colonies was around 12 mU/mg, while in the BBMV preparations it was around 70 mU/mg, indicating an enrichment of APN activity of around 5.8-fold. Importantly, no significant differences (P>0.05) in APN activity were observed between the Vip-Unsel and Vip-Sel colonies. For the Vip-Unsel colony, specific ALP activity was 7.44 mU/mg in midgut homogenates and 42.5 mU/mg in the BBMV, with an enrichment of 5.8-fold for the BBMV preparation, in agreement with the value obtained for APN. In contrast, dramatically reduced ALP activity was detected in samples from the Vip-Sel colony (1.15 mU/mg in midgut homogenate and 1.88 mU/mg in BBMV). While unexpected, this observation is in line with reports of reduced ALP levels in Cry1-resistant lepidopteran species, including *H. virescens* (20-23). Consequently, we further explored the extremely reduced ALP activity in Vip-Sel to determine whether it was due to a loss of function or reduced gene expression.

**Figure 2.**
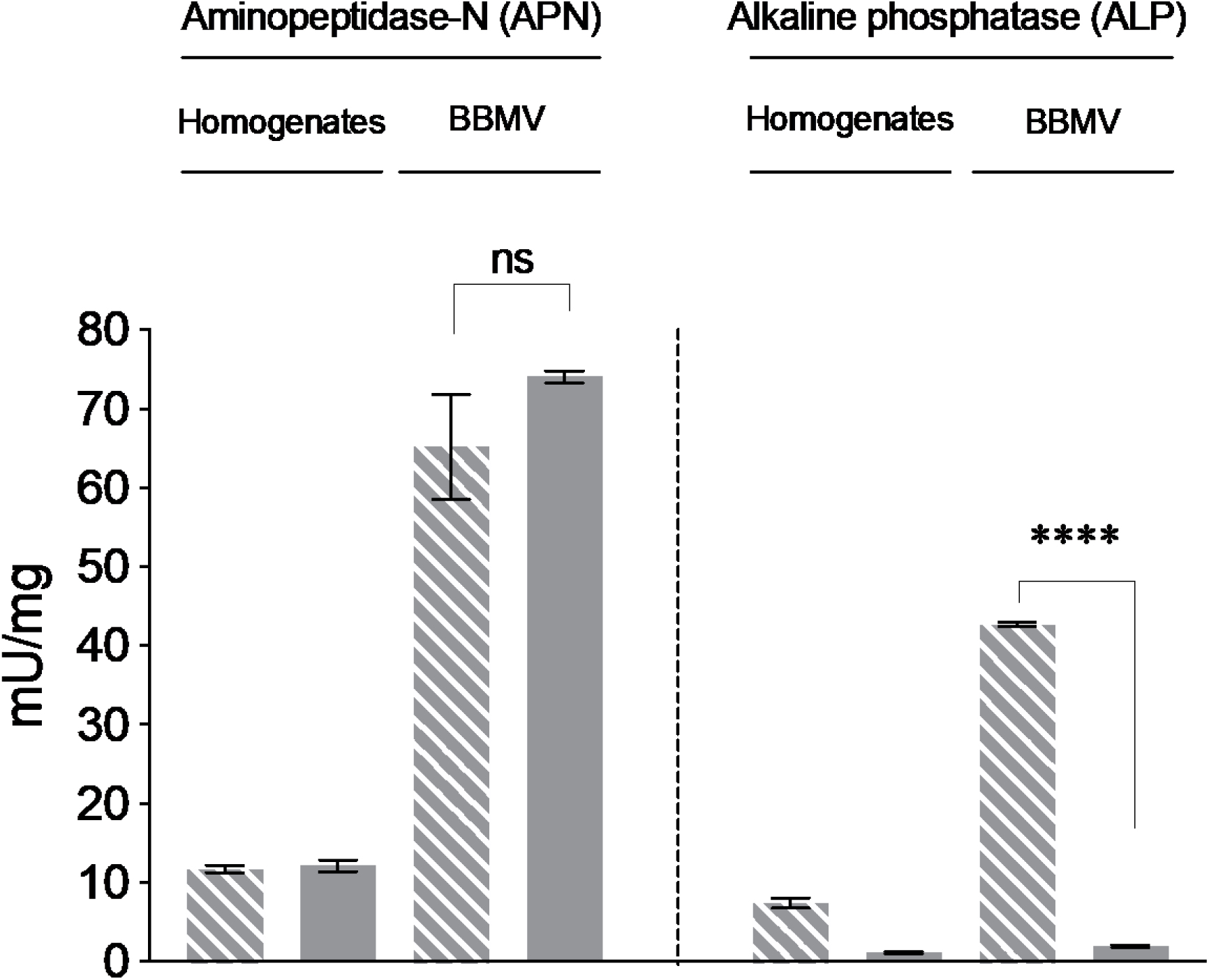
Enzymatic activities in homogenates and BBMV from the two colonies of *H. virescens*. (Dashed-grey bars: Vip-Unsel, grey bars: Vip-Sel). Each bar represents the mean of three replicates (± SEM). Asterisks represent significant difference (Student’s *t* test, *P* < 0.0001).

Electrophoretic comparison of BBMV proteins from the two *H. virescens* colonies showed a protein band of ∼66 kDa for the Vip-Unsel colony that was almost imperceptible in the BBMV from the Vip-Sel colony (Figure 3, panel A). Western blotting indicated the presence of ALP in the ∼66 kDa protein band, and confirmed the highly reduced levels of this protein in the Vip-Sel colony (Figure 3, panel B). The composition of the ∼66 kDa protein band and its relative abundance in the two *H. virescens* colonies were determined by liquid chromatography coupled to mass spectrometry (LC-MS) analysis. The spectra for the most abundant protein detected and identified in the ∼66 kDa band matched to membrane-bound alkaline phosphatase (mALP) from *H. virescens* (Genbank Accesion no. ABR88230). According to the exponentially modified protein abundance index (emPAI) expressing the proportional protein content in a protein mixture, the abundance ratio of mALP between Vip-Unsel and Vip-Sel was 22.7-fold.

**Figure 3.**
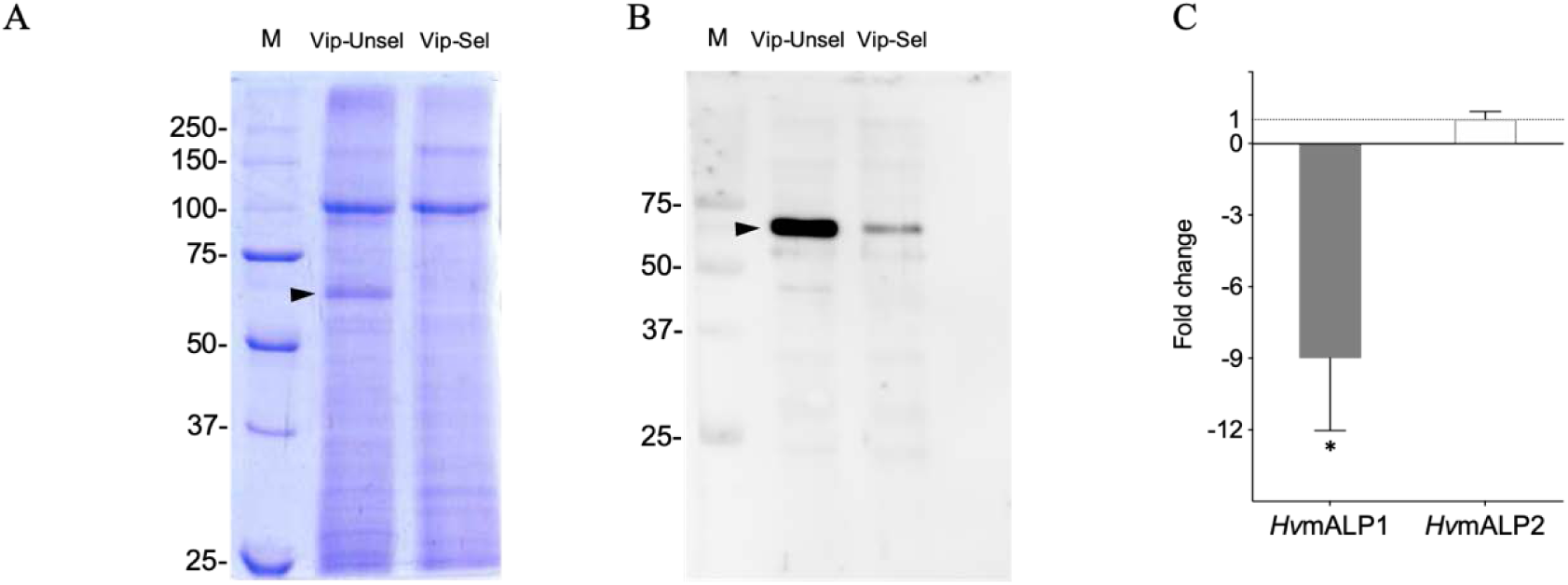
Analysis of membrane ALP levels in the susceptible (Vip-Unsel) and resistant (Vip-Sel) colonies of *H. virescens*. **A)** Protein gel electrophoresis (SDS-PAGE) of BBMV from the two colonies. **B)** Western blot performed with anti-ALP antibody against BBMV from the two colonies. Black arrow indicates mALP (*ca.* 66 kDa). Lane M: Protein marker (molecular weight in kDa). **C)** Membrane ALP expression levels in Vip-Sel colony using transcript levels in Vip-Unsel colony as a reference. Fold-changes calculated by REST-MCS Software. Bars represent the mean of three independent experiments (± SD, * *P* < 0.05).

To test if the reduced mALP protein levels in Vip-Sel were controlled at the transcriptional level, we performed real time quantitative PCR (RT-qPCR) with total RNA from the two colonies. Transcript levels for two *H. virescens* mALP genes, *HvmALP1* (accession number FJ416470.1) and *HvmALP2* (accession number FJ416471.1), were analyzed. Compared to insects from the Vip-Unsel colony, larvae from the Vip-Sel colony had significant (P-value < 0.05) 9-fold down-regulation of the *HvmALP1* gene, while transcript levels for *HvmALP2* were not different between colonies (Figure 3, panel C). These results support that reduced ALP enzyme activity in BBMV from Vip-Sel compared to Vip-Unsel is due to reduced expression of *HvmALP1* in the Vip-Sel colony.

### Ligand blotting

Since *H. virescens* ALP was proposed to play a role in binding of Cry1 proteins to the midgut membrane (24), we used ligand blotting to test whether mALP was involved in Vip3Aa binding. Binding of Vip3Aa to blots of resolved BBMV proteins was detected with anti-Vip3Aa antisera. No differences in the Vip3Aa-binding band pattern were detected between both colonies, in agreement with the binding results with radiolabeled Vip3Aa. However, no Vip3Aa binding was observed at the mALP position (∼ 66 kDa) (Figure 1, panel B).

### Transient expression of ALP in cells

To further test the potential role of mALP as functional Vip3Aa receptor, we cloned and transiently expressed the *HvmALP* gene in cultured (Sf21) insect cells and performed cell viability tests after challenge with Vip3Aa. Transfection was successful, as transfected cells showed ∼5-fold increased specific ALP activity compared with non-transfected cells or cells transfected with the empty plasmid (Figure 4, panel A). However, after a challenge with Vip3Aa, the viability of transfected cells was not significantly different (P > 0.05) from that of the control cells (Figure 4, panel B), confirming that mALP does not serve as functional receptor for Vip3Aa during the toxicity process.

**Figure 4.**
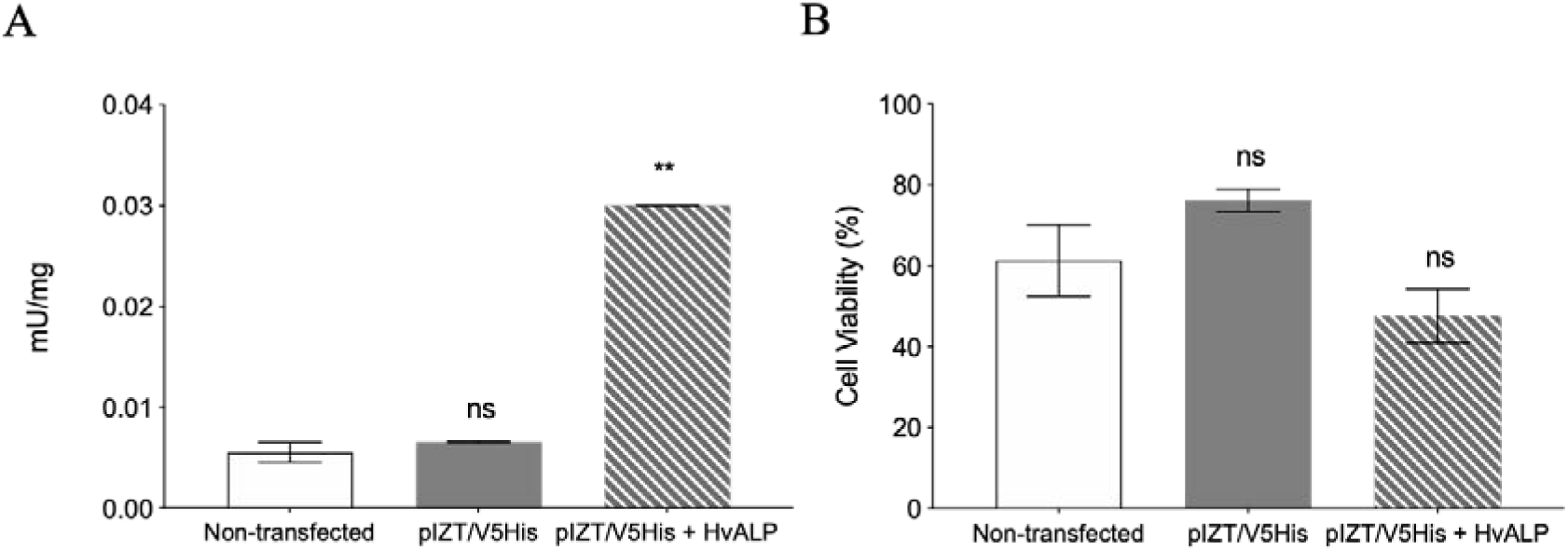
Specific ALP enzymatic activity and viability assays of Sf21 cells producing the HvmALP1 isoform. **A)** Alkaline phosphatase enzymatic activity on non-transfected cells (empty bars), cells transfected with empty plasmid (grey bars) and plasmid with *HvmALP1* (dashed-grey bars). **B)** Cell viability after 24 hours of Vip3Aa intoxication on the same three cell types. Each value represents the mean (± SEM). Means were compared by Student’s *t*-test (** *P* < 0.01).

## DISCUSSION

The use of resistant insect strains isolated from the field or selected in the laboratory has been a powerful tool to understand the biochemical and genetic bases of resistance to Bt insecticidal proteins. Many studies have found that the alteration of the membrane receptors is a common mechanism conferring high levels of resistance to Cry proteins (25-27). In the case of Cry1 proteins, an important body of literature identifies the main receptors altered in association with resistance, including aminopeptidase N, ABC transporters, cadherins and membrane alkaline phosphatases (28,29). In contrast, three candidate receptors have been proposed for Vip3A proteins, including the *Spodoptera spp.* ribosomal protein S2 (30), the fibroblast growth factor receptor-like protein (31) and the scavenger receptor class C-like protein (32). However, the relationship between these proposed receptors and Vip3Aa resistance has not been established.

In the present work, we aimed to determine whether alteration of membrane receptors was the basis for the observed 2,040-fold resistance to Vip3Aa in a *H. virescens* laboratory colony. Results from binding assays with BBMV and radiolabeled Vip3Aa did not detect significant difference between the susceptible and resistant colonies, suggesting no involvement of binding site alteration in resistance. This conclusion was further supported by results from ligand blotting, where no differences between the binding patterns of Vip3Aa to BBMV proteins from the two colonies was observed. Similar results were reported for a laboratory-selected Vip3A-resistant colony of *H. armigera* (17), suggesting that high levels and narrow spectrum Vip3A resistance may develop by mechanisms other than alteration of Vip3Aa binding sites.

Despite the fact that differences in binding were not found between the two *H. virescens* colonies, a dramatic reduction in the ALP enzymatic activity was detected in midgut samples from the resistant compared to susceptible colony. Western blot and RT-qPCR analyses showed that the decreased activity was due to a reduction in the amount of mALP protein, which was controlled at the transcriptional level. Importantly, only one of the ALP isoforms (*HvmALP1*) appeared down-regulated, while transcript levels for the *HvmALP2* isoform were not different between colonies. Taken together, these results support the hypothesis that reduced expression of the *HvmALP1* isoform is associated with Vip3Aa resistance in the Vip-Sel colony. Down-regulation or reduced levels of mALP in the midgut membrane have been observed as a common phenomenon in several Cry-resistant insects such as resistance to Cry1Ac in *H. virescens* (24), *Helicoverpa zea* (20), *Plutella xylostella* (21), and *Helicoverpa armigera* (23), to Cry1F in *S. frugiperda* (22), to Cry1C in *Spodoptera litura* (33), and even in *Aedes aegypti* resistant to a Bt susbp. *israeliensis* (Bti) (34). The fact that Cry1Ac and Cry1C have different binding sites (35) suggests that the role of ALP down-regulation in resistance may not be related to reduced Cry binding. In agreement with this hypothesis, reduced ALP levels were associated with resistance but Vip3Aa binding was not detected in Vip-Sel. In addition, susceptibility of Sf21 cells expressing HvmALP1 was not significantly different to Vip3Aa, supporting that ALP is not a functional receptor for Vip3Aa in *H. virescens*. However, gene silencing experiments provide evidence that ALPs are somehow involved in the intoxication process of Cry1, Cry2 and Cry11 proteins in several insect species (36-41).

In a Cry1Ac-resistant colony of *P. xylostella*, altered expression of different genes (including the *Px*mALP) was reported to be *trans*-regulated by up-regulation of a mitogen-activated protein kinase, which was linked to resistance (21). Similar *trans*-regulation of genes involved in resistance to Bt has also been observed for APN in *Trichoplusia ni* resistant to Cry1Ac (42) and *Ostrinia furnacalis* resistant to Cry1Ab (43), and for both APN and an ABCC transporter in *Bombyx mori* resistant to Cry1Ab (44). Further research should test the involvement of this control mechanism in down-regulation of *mALP* in Bt-resistant colonies.

The data in this study provides for the first time evidence that reduced levels of mALP are associated with Vip3Aa resistance. Considering that orthologues of this *mALP* are commonly down-regulated in Cry1-resistant lepidopteran pests (23), our results suggest that the potential use of *mALP* down-regulation as resistance marker may extend to resistance against Vip3A proteins.

## MATERIALS AND METHODS

### Insects

Two colonies of *H. virescens* originated from the same field population collected in Arkansas (USA) were used in this study: Vip-Sel (Vip3Aa-resistant) and Vip-Unsel (Vip3Aa susceptible). The process of selection of the Vip-Sel colony with Vip3Aa has been previously described (12,18). After 13 generations of selection, the LC_50_ of the Vip-Sel colony was 2,300 μg ml^-1^, representing a 2,040-fold resistance ratio relative to the control Vip-Unsel colony. Both colonies were reared at the Imperial College London, Silwood Park campus (UK), and frozen larvae were sent for analysis to the University of Valencia (Spain).

### BBMV preparation and enzyme activity assays

Brush border membrane vesicles (BBMV) from 3^rd^ instar *H. virescens* larval midguts from Vip-Sel and Vip-Unsel colonies were prepared according to the differential magnesium precipitation method (45). Isolated BBMV were flash frozen in liquid nitrogen and kept at -80°C until used. The protein concentration of the BBMV preparations was determined by the method of Bradford using bovine serum albumin (BSA) as a standard (46).

Alkaline phosphatase (ALP) and leucine aminopeptidase (APN) activities were used as brush border membrane enzymatic markers to determine the quality of the BBMV preparations (47). Specific ALP activity was determined by chromogenic detection of *p*-nitrophenyl phosphate (PNPP) substrate hydrolysis into *p*-nitrophenol, and specific APN activity was detected by hydrolysis of L-Leu-*p*-nitroanilide substrate into *p*-nitroanilide. In both cases, chromogenic variation was measured on 1 μg of either BBMV or midgut homogenate at 405 nm (Infinite m200, Tecan, Mannedorf, Switzerland). Two different batches of BBMV were used and all enzymatic activity assays were performed in triplicates. Means values for enzyme activities from Vip-Unsel and Vip-Sel were compared by Student’s *t*-test at 5% level of significance.

For measuring specific ALP enzymatic activity in cultured Sf21 cells, a 1.6 ml suspension of each cell type (non-transfected, transfected with empty plasmid, and transfected with ALP) was used. Culture cells were centrifuged, washed twice with 300 μl of phosphate buffered saline (PBS) and then resuspended in 50 μl of PBS. Protein concentration was determined by the method of Bradford and specific ALP activity measured as above.

### Vip3Aa protein expression and purification

The Vip3Aa16 (Vip3Aa) protein (NCBI accession No. AAW65132) was overexpressed in recombinant *Escherichia coli* BL21 carrying the *vip3Aa16* gene. Protein expression and lysis was carried out following the conditions described elsewhere (48). Soluble Vip3Aa in the cell lysate was purified by two different methodologies. For binding and cell viability assays, Vip3Aa was partially purified by isoelectric point precipitation (IPP), activated with trypsin treatment and further purified by anion-exchange chromatography, as previously described (11). For ligand assays, affinity chromatography purification was carried out using a HiTrap chelating HP column (GE Healthcare) and then activated with trypsin, as described (11).

### Vip3Aa labeling and binding experiments

Purified Vip3Aa activated protein (25 μg) was labeled with 0.5 mCi of ^125^I using the chloramine T method (11). The labeled protein was separated from the excess of free ^125^I in a PD10 desalting column (GE Healthcare Life Sciences, United Kingdom) and the purity of the ^125^I-labeled Vip3Aa was checked by autoradiography. The specific activity of the labeled protein was 2.2 mCi/mg.

Binding assays were performed as described elsewhere (11). Prior to being used, BBMV were centrifuged and resuspended in binding buffer (20 mM Tris, 150 mM NaCl, 1 mM MnCl_2_, pH 7.4, 0,04% Blocking reagent from Sigma Aldrich). Competition binding experiments were conducted by incubating 1.4 μg of BBMV protein with 0.65 nM ^125^I-Vip3Aa in a final volume of 0.1 ml of binding buffer for 90 min at 25°C in the presence of increasing amounts of unlabeled Vip3Aa. After incubation, samples were centrifuged at 16,000 x *g* for 10 min and the pellet was washed once with 500 μl of cold binding buffer. Radioactivity retained in the pellet was measured in a model 2480 WIZARD^2^ gamma counter. Data from the competition experiments were analyzed to determine equilibrium binding parameters, dissociation constant (*K*_d_) and concentration of binding sites (*R*_t_) using the LIGAND software (49).

### Western and ligand blotting

For the detection of ALP proteins in BBMV by Western blotting, BBMV (20 μg) were suspended in ice-cold PBS and heat denatured before separation on a SDS-10% PAGE gel. The resolved BBMV proteins were transferred to a nitrocellulose filter (Protran 0.45 μm NC, GE Healthcare) using a BioRad Mini Trans-Blot system at 4°C in blotting buffer (39 mM Glycine, 48 mM Tris-HCl, 0.037% SDS, 10% methanol, pH 8.5) for 1 h at constant voltage (100 V). After transfer, the nitrocellulose filter was blocked in blocking buffer (PBS, 0.1% Tween 20, 5% skimmed milk powder) overnight at 4°C. After blocking and washing with PBST (PBS, 0.1% Tween 20) three times (5 min each), incubation with primary antibody against the membrane-bound form of ALP from *Anopheles gambiae* (generously provided by Dr. M. Adang, University of Georgia, USA) was performed for 90 min at a 1:5,000 dilution at room temperature (RT). The membrane was then washed with PBST three times for 5 min each and then incubated with secondary antibody (goat anti-rabbit conjugated to horseradish peroxidase [HRP] at a 1:10,000 dilution) for 1h at RT. After being washed with PBST three times for 5 min each, the membrane was developed using enhanced chemiluminescence (ECL Prime Western Blotting detection reagent, GE Healthcare) in an ImageQuant LAS 4000 (GE Healthcare), according to the manufacturer’s instructions.

Ligand blotting for the detection of BBMV proteins binding Vip3Aa protein was performed with BBMV proteins resolved and immobilized as described above for Western blotting. The nitrocellulose membrane was blocked for 1 h at 4 °C in blocking buffer (5% skimmed milk), and after three washes for 5 min each with PBST buffer, it was incubated overnight at 4 °C with blocking buffer (1% skimmed milk) supplemented with affinity chromatography-purified Vip3Aa at a final concentration of 4 μg/ml. After washing with PBST three times for 5 min each, the membrane was incubated with primary antibody against Vip3Aa at a 1:5,000 dilution for 1 h at RT. After three washing steps with PBST (5 min each), membranes were incubated with secondary antibody (goat anti-rabbit conjugated to HRP) for 1 h at RT. Upon washing three times (5 min each) with PBST, the membrane was developed as described for Western blotting.

### Proteomic analysis

After resolving BBMV proteins from Vip-Sel and Vip-Unsel colonies by SDS-10% PAGE, the gel was stained with Coomassie blue (Thermo Scientific™). The band corresponding to the expected molecular weight of ALP (∼ 66 kDa) was cut out and subjected to analysis by nano-electron spray ionization (nano-ESI) followed by tandem mass spectrometry (qQTOF) in a 5600 TripleTOF (ABSCIEX) system. Results were analyzed with ProteinPilot v5.0 software and the relative amount of the proteins detected was estimated using the exponentially modified protein abundance index (emPAI) as described elsewhere (50).

### RT-qPCR

Relative expression levels for *HvmALP1* and *HvmALP2* isoforms (accession numbers FJ416470 and FJ416471, respectively) were determined by reverse transcription quantitative polymerase-chain reaction (RT-qPCR). For this purpose, total RNA of dissected midguts from both colonies (Vip-Unsel and Vip-Sel) was isolated using RNAzol (MRC Inc., Cincinnati, OH) according to the manufacturer’s protocol. Each RNA (1 μg) was reverse-transcribed to cDNA using random hexamers and oligo (dT) by following the instructions provided in the Prime-Script RT Reagent Kit (Perfect Real Time from TaKaRa Bio Inc., Otsu Shiga, Japan). RT-qPCR was carried out in a StepOnePlus Real-Time PCR system (Applied Biosystems, Foster City, CA). Reactions were performed using 5× HOT FIREpol EVAGreen qPCR Mix Plus (ROX) from Solis BioDyne (Tartu, Estonia) in a total reaction volume of 25 μl. Specific primers for *HvmALP1, HvmALP2* and *Rps18* (endogenous control) genes were as described elsewhere (23). The REST MCS software was used for gene expression analysis (51).

### Expression vector construction

The full-length *HvmALP1* transcript was amplified from cDNA of *H. virescens* larvae and cloned into pET30a as described elsewhere (52). Purified plasmid DNA was digested with *Eco*RI and *Not*I to excise the full-length sequence and ligate it in frame into *Eco*RI-*Not*I sites of the pIZT/V5His vector (Invitrogen), to generate the pIZT/V5His/*HvmALP1* construct. Ligation products were transformed into *E. coli* strain DH5α and transformants checked for correct insertion by sequencing (University of Tennessee Sequencing Facility). Purified plasmid was used to transform *E. coli* strain DH10β and liquid cultures of LB medium supplemented with Zeocin (25μg/ml) were used to amplify the vector. To purify the plasmids for transfection, the NucleoSpin® Plasmid kit (Macherey-Nagel, Düren, Germany) was used. Double digestion with *Eco*RI and *Not*I (which cleaved the full length *HvmALP1* insert) and 1% agarose gel electrophoresis were performed to check plasmid and/or insert integrity. The concentration of plasmid DNA was measured with a Thermo Scientific™ Nanodrop™ 2000 Spectrophotometer.

### Transient expression of *HvmALP1* in Sf21 cells

Cultured Sf21 insect cells, originally derived from *S. frugiperda*, were maintained in 25 cm^2^ tissue culture flasks (T25 flasks, Nunc) at 25°C with 4 ml of Gibco® Grace’s Medium (1x) (Life Technologies™, Paisley, UK) supplemented with 10% heat-inactivated fetal bovine serum (FBS).

For transient expression, cells were seeded on 12-well plates with the same medium without FBS at *ca*. 70% confluency and transfected with 0.5 μg of the pIZT/V5His/*HvmALP1* or pIZT/V5His plasmid using Cellfectin® II Reagent (Invitrogen), following manufacturer’s instructions. Five hours post-transfection, the medium was replaced with fresh medium containing 10% FBS. After 24 hours, cells were examined using a confocal microscope (Olympus, FV1000MPE, Tokyo, Japan) equipped with the appropriate filter for green fluorescent protein (GFP) detection as transfection marker. The enzymatic activity of alkaline phosphatase was then measured as explained above.

### Cell viability assays

Viability of transfected Sf21 cells exposed for 24 hours to Vip3Aa was measured using the MTT (3-[4,5-dimethylthiazol-2-yl]-2,5-diphenyltetrazolium bromide) assay. Briefly, cells (100 µl per well) were transferred to 96-well ELISA plates (flat bottom) and incubated at 25°C for at least 45 min. Then, 10 µl of trypsin-activated Vip3Aa toxin was added to each well at a final concentration of 300 µg/ml. As negative and positive controls, 10 µl of Tris buffer (Tris 20 mM, NaCl 150 mM, pH 9) and 10 µl of 2% Triton X-100 were used, respectively. After 24 hours of incubation at 25°C, cell viability was assessed by applying 20 µl of CellTiter 96® AQueous One Solution Reagent (Promega, Madison WI) to each well and incubating for 2 h at 25°C. Absorbance was measured at 490 nm (Infinite m200, Tecan, Mannedorf, Switzerland). The percentage of viable cells was obtained as described elsewhere (53). Mean values in the transfected cells against the non-transfected cells were compared by Student’s *t*-test at 5% level of significance.

## ACKNOWLEDGMENTS

This research was supported by the Spanish Ministry of Science, Innovation and Universities (grant no. RTI2018-095204-B-C21). D.P. is recipient of a PhD grant from the Spanish Ministry of Science, Innovation and Universities (grant ref. FPU15/05652). Support for JLJ-F was provided by the NC246 Multistate Hatch project from the USDA National Institute for Food and Agriculture (NIFA). The proteomic analysis was performed at the Proteomics Facility of the Servei Central de Suport a la Investigació Experimental (SCSIE) at the Universitat de València (Valencia, Spain) that belongs to ProteoRed, PRB2-ISCII, supported by grant PT13/0001, of the PE I+D+I 2013-2016, funded by ISCII and FEDERPT13/0001.

